# Genetic dependencies associated with transcription factor activities in human cancer cell lines

**DOI:** 10.1101/2023.02.23.529701

**Authors:** Venu Thatikonda, Verena Supper, Madhwesh C. Ravichandran, Jesse J. Lipp, Andrew S. Boghossian, Matthew G. Rees, Melissa M. Ronan, Jennifer A. Roth, Sara Grosche, Ralph A. Neumüller, Barbara Mair, Federico Mauri, Alexandra Popa

## Abstract

Transcription factors (TFs) are key components of the aberrant transcriptional programs in cancer cells. In this study, we used TF activity (TFa), inferred from the downstream regulons as a potential biomarker to identify associated genetic vulnerabilities in cancer cells. Our linear model framework, integrating TFa and genome-wide CRISPR knockout datasets identified 1,770 candidate TFa-target pairs across different cancer types and assessed their survival impact in patient data. As a proof of concept, through inhibitor screens and genetic depletion assays in cell lines, we validated the dependency of cell lines on predicted targets linked to TEAD1, the most prominent TF from our analysis. Overall, these candidate pairs represent an attractive resource for early-stage targets and drug discovery programs in oncology.

## Introduction

Transcription factors (TFs) bind regulatory elements such as enhancers or promoters in order to regulate the expression of genes. The role of TFs has been well established in health and disease, including human cancers^1^. TF activities are reprogrammed in many different cancers due to genetic alterations such as amplifications (e.g., *NMYC, CMYC)*^2^ and epigenetic mechanisms including the formation of *de novo* enhancers (e.g., FOXA1, MITF)^3^. Certain TFs sustain oncogenic transcriptional programs required for tumor maintenance that differ from those in neighboring normal cells. In addition, abnormal activities of TFs modulate hallmark properties of cancers such as the adaptive response of cancer cells to therapy. This is achieved through the regulation of processes such as epithelial to mesenchymal transition (EMT) and acquired resistance to chemotherapy or targeted therapy. For example, RUNX2 drives EMT in breast cancer and prostate cancers and mediates chemoresistance in melanoma and TEA domain transcription factors (TEAD) TFs regulate phenotypic plasticity ^4–8,9^. Given the central role of TFs in tumor biology, their selective pharmacological inhibition constitutes an attractive therapeutic strategy to treat human malignancies. Developing small molecule inhibitors for cancer relevant proteins such as TFs is challenging and often hampered by the absence of binding pockets or structural enablement. While there have been some recent advances in drug development for challenging targets through methods such as PROTACs^10^, or leveraging synthetic lethality^11^, there is still a need to expand the range of targets in cancer cells. Identifying additional partner proteins or dependencies associated with TF function in tumor cells could provide new opportunities for targeted therapies.

Genome-scale loss-of-function (LOF) studies conducted in hundreds of human cancer cell lines using CRISPR/Cas9 and shRNA-based strategies have successfully identified tissue and context specific genetic dependencies across tumor types^12^,^13–15^. Integrating genome-wide essentiality data with complementary multi-omics data enables the identification of potential biomarkers that are predictive of specific gene dependencies. Theseanalyses, for example, revealed that micro-satellite unstabletumors are particularly vulnerable to WRN LOF, as well as, that PRMT5 constitutes a potential therapeutic target in *MTAP* deleted cancers.^16–21^ In addition, concepts such as synthetic lethality have also been applied within the paralogue gene space to identify potential biomarkers (e.g., a genetic alteration) and target gene pairs. For instance, mutations in STAG2 are predictive of sensitivity to *STAG1* LOF^22–25^, and loss of *DDX3Y* (through the loss of chromosome Y) is predictive of sensitivity to DDX3X LOF^26^.

Here we provide a framework for the systematic identification of gene dependencies associated with TF activities across different tumor types. To this end, we integrated three genome-scale LOF studies (Project Achilles^27,28^, Project Sanger^29^, Project Drive^30^) with transcriptomic data from 859 human cancer cell lines spanning 27 tumor types, including 757 solid cancers and 102 hematological cancer cell lines. We extensively annotated candidate TF activity (TFa) - gene dependency associations in terms of their tumor type specificity and clinical relevance using The Cancer Genome Atlas^31^ (TCGA) patient dataset. Finally, we observed TEAD1 TFa to be associated with a large number of gene dependencies and further characterized these associations through inhibitor screens and single gene *in vitro* CRISPR-Cas9 depletion assays. Thus, providing proof of the potential benefits ensured by our approach in the identification of new surrogate targets of oncogenic TFs.

## Results

### Identification of pan-cancer TF activity-associated gene dependencies

We identified associations between a TF activity biomarker (TFa) and gene dependencies (target), hereafter termed “TFa-target pair” using expression and dependency data in cell lines. Since TFs regulate the expression of downstream genes, also termed regulons, we defined the combinatorial expression of annotated TF regulons as the TF activity for each cell line ^32^ (see methods). We removed all protein coding genes (including TFs) which are potentially essential in cancer and normal cells (pan-essential), as well as genes that are not essential for cell fitness (never essential) based on dependency data. We then implemented a linear model-based framework on 92 context-essential TFs (Supp Fig 1a, see methods) to identify their activity associated gene dependencies. In the linear model, we used TFa as the independent variable and gene CRISPR knock-out (KO) dependency score as the dependency variable and the tumor type of origin of the cell line as the covariate (Fig 1a). The inclusion of cell line tumor type in the model accounts for the variability in dependency score due to tissue type differences. We applied this framework across 757 solid cancer cell lines in Project Achilles^27,28^ and tested over a million linear models. We performed TF-specific false discovery rate (FDR) adjustment and considered the associations with FDR <10% and an effect size (TFa coefficient) of greater than 3x the standard deviation as significant TFa-target pairs. To increase the robustness of our results, we repeated the linear modeling framework with two additional genome-wide dependency datasets: CRISPR KO data with 304 solid cancer cell lines (Project Sanger^29^) and shRNA data with 635 solid cancer cell lines (Project Drive^30^, Supp Table 1). The final set of TFa-target pairs consisted of pairs that were identified as significant in at least two out of three dependency databases(Project Achilles, Sanger, Project Drive). We performed downstream analysis of on the final set of pairs to assess their impact on survival using TCGA data^31^, and validated three gene dependencies associated with TEAD1 TFa using CRISPR single gene KO experiments (Fig 1a).

**Figure 1:**
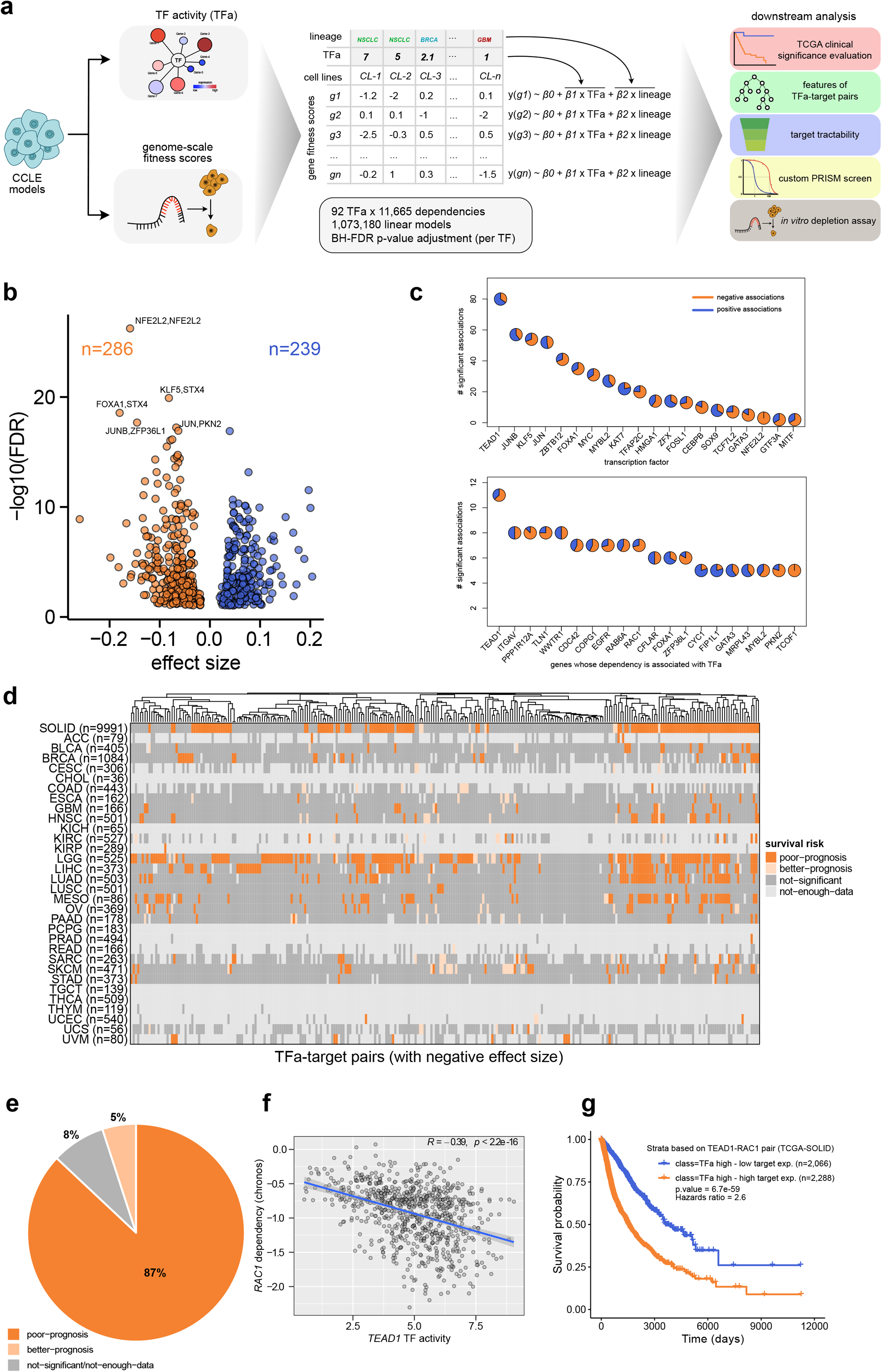
Framework and overview of the TFa-gene dependency associations. a) Linear model based framework to identify target gene dependencies associated with transcription factor activities (TFa). Further downstream analysis of the significant TFa-target pairs performed in this study are indicated. b) Volcano plot of TFa-target pairs identified from our pipeline using 757 solid cancer cell lines. Top associations are labeled. Negative effect size indicates increased dependency associated with increased TFa and positive effect size indicates increased dependency associated with decreased TFa. c) Top panel: TFs with the highest number of total associations as TFa biomarker, bottom panel: dependency targets with the highest number of significant associations as dependency feature. d) Heatmap illustrating the impact of a TFa-target pair on survival of different TCGA cancer types when both TFa activity and associated target expression are high. Analysis details are illustrated in Supp Fig 1a. Dark orange representsnegative pairs with poor prognosis in TCGA. Light orange represents negative pairs with better prognosis in TCGA. Dark grey and light grey are negative pairs without a TCGA statistically significant survival impact or too few patient samples, respectively. e) Distribution of fraction of TFa-target pairs and their association with survival. f) Scatter plot showing the correlation of TEAD1 TF activity and RAC1 dependency. g) Kaplan-Meier plot showing the survival difference between RAC1 high vs low expression subgroups within TEAD1 TFa activity high tumors.

Overall, we identified 525 TFa-target candidate pairs for which TFa was significantly associated with target gene dependency, comprising 26 TFs as TFa biomarkers and 226 gene dependencies as potential targets (Supp Table 2). The candidate pairs include both negative and positive associations (Fig 1b) where negative associations (n=286) represent increased dependency (*i*.*e*., lower dependencyscore) of a given gene with increased TFa and positive associations (n=239) represent increased dependency with decreased TFa(Fig 1b). Among the predicted associations, we identified seven self-dependent TFs *i*.*e*., increased dependency of a TF was associated with its own increased activity (e.g., NRF2/NFE2L2, JUN), previously reported synthetic lethal associations (e.g., GATA3-MDM2^33^) and previously unknown associations (e.g., TEAD1-RAC1, KLF5-STX4). We also observed TFa-target pairs where the associated dependency target gene is also a TF (e.g., TEAD1-FOSL1), suggesting that essentiality of a TF can be potentially predicted by the activity of associated TFs (Supp Table 2). Among the TFs with significant associations in our final list of candidate pairs, we observed TEAD1 to have the largest number of associations (Fig 1c) both as a TFa biomarker as well as a potential dependency target. Identification of TEAD1 in both negative and positive interactions is in line with recent studies emphasizing the dual role of the YAP/TEAD axis across solid tumors, with both anti- and pro-cancer effects^34^ (Fig 1c).

Next, we investigated the potential impact of the identified TFa-target pairs on patient survival using TCGA expression and clinical data. We only considered the 286 TFa-target candidate pairs with negative effect size, since (Fig 1b) increased activity of TFs is potential evidence of involvement in oncogenic transcriptional programs. To this end, we classified all TCGA tumors into subgroups based on TFa and its associated target gene expression (Supp Fig 1b; see methods). We compared the survival difference between high and low target gene expression subgroups within high TFa tumors using a Cox proportional hazards model^35^ (Supp Fig 1b). We reasoned that low expression of a target gene in the presence of high TFa (TFa high-low target exp.) would have better prognosis compared to a high expression subgroup (TFa high-high target exp.), in line with our hypothesis. This analysis was conducted separately for each cancer type in the TCGA dataset and pan-cancer, combining all solid tumors. Of the 286 TFa-target candidate pairs, 251 pairs (∼87%) were associated with poor patient survival in at least one clinical context, when TFa and its associated gene expression were both high (TFa high-target high) (Fig 1d,e). Among these pairs, as an example, we show TEAD1-RAC1, where TEAD1 TFa is strongly associated with RAC1 dependency across cell lines (Fig 1f), and the patient subgroup with TEAD1 high TFa and RAC1 high expression (within solid tumors) showed poor survival compared to high TEAD1 TFa and low RAC1 expression patient subgroup (Fig 1g, Supp Table 3).

We independently applied the same analysis framework to 102 hematological cancer cell lines in Project Achilles. In total we identified 101 TFa-target pairs with negative effect size in hematological cancers (Supp Fig 1c, Supp Table 4). We observed an association with poor prognosis in 74/101 (∼74%) of these pairs when TFa and its associated gene expression are high in TCGA hematological malignancies (Supp Fig 1c, d, e, Supp Table 5). We then tested the overlap of TFa-target pairs identified across solid and hematological cancer cell lines and observed no pairs in common, illustrating the molecular differences of these cancer categories^36^ (Supp Fig 1d). We further tested the overlap between the dependency targets identified from our framework for each TF and their annotated regulons. We observed a very low Jaccard score (average Jaccard score of 0.005 across all TFs), a metric indicating the extent of overlap ranging from 0 (no) to 1 (full overlap) (Supp Fig 1f, Supp Table 6), suggesting the dependency targets identified through our framework are complementary to the annotated regulon of each TF.

In summary, we identified 286 TFa-target candidate pairs across solid cancer cell lines showing increased dependency of target gene with increased TFa. We have shown that 87% of these candidate pairs are associated with poor survival in patients with both high TFa and target expression in at least one tumor type. While our in-depth analysis is focused on solid tumor types, our proposed framework can be further applied to other tumor subtypes, potentially leading to the identification of further candidate pairs.

### Features of TFa-target pairs

We sought to understand the determinants of TFa-target synthetic lethality relationship, i.e. increased dependency of target gene association with increased TFa. To this end, we explored 272 candidate TFa-target pairs (excluding self-dependent TFs and pairs where the TFa and target are both TFs) termed ‘significant pairs’. As a control we prepared a ‘non-significant’ random set of 300 pairs that were not significant but had a negative effect size from our initial linear modeling analysis. For this combined set of 572 pairs, we collected 15 different features that capture at various levels how the gene pairs are associated e.g., protein-protein interaction (PPI), expression correlation, genetic dependency correlation or pathway overlap (TF and target both are in the same molecular pathway; Supp Table 7)^37^.

In order to assess each feature’s power in identifying TFa-target significant relationships, we treated each feature independently as a classifier and computed the area under the receiver operating curve (ROC AUC) (Fig 2a). The top predictive feature was the direct interactors overlap between TF and associated target (ROC AUC = 0.73). This feature indicates that “significant pairs” shared more direct interactors in the protein-protein interaction (PPI) network than the control pairs (Fig 2b). Next, we wanted to determine if there was a significant overlap of these direct interactors between TF and its associated target gene. We observed that significant pairs show a stronger p-value for overlap (Fisher’s exact test, Supp Fig. 2a). Similarly, we observed a higher Jaccard score of overlap for significant pairs, a metric indicating the extent of overlap (Supp Fig. 2a). A previous analysis on the identification of synthetic lethal pairs within paralog families revealed that the higher the number of common interactors between two genes in a protein-protein interaction network, the higher the chance for these two genes to be functionally redundant and part of a synthetic lethal genetic interaction^37^. In line with these findings, our results suggest that a high number of common interactors between a TF and its associated target gene is a predictor of a putative TFa-target significant association. Similarly, we observed a strong overlap of pathways (ROC AUC 0.71) and dependency correlations of a TF and its associated target (ROC AUC 0.67) for significant pairs compared to control non-significant pairs (Fig 2b). The second strongest feature with an ROC AUC of 0.71, “pathway overlap” indicates the number of common pathways in which TF and associated target gene are involved (Fig. 2a). Similar to the feature “direct interactors overlap” described here, the number of common pathways in which TF and associated target gene are involved is significantly higher for significant pairs compared to non-significant pairs (Fig 2b). Finally, we observed significant pairs show a strong positive correlation TF and its associated target gene dependencies (Fig. 2b), suggesting the co-essentiality of these pairs in cancer cells. A recent study^38^ employed the correlation of essentialities from genome wide CRISPR KO screens to identify genes with similar molecular functions. Consistent with these studies, our results suggested that our candidate TFa-target pairs could be engaging in similar biological mechanisms. In addition to these features, we also observed statistically significant differences between significant and non-significant pairs for features such as “TF and its target gene expression correlation”, “direct interaction evidence” (Supp Fig 2a, b).

**Figure 2:**
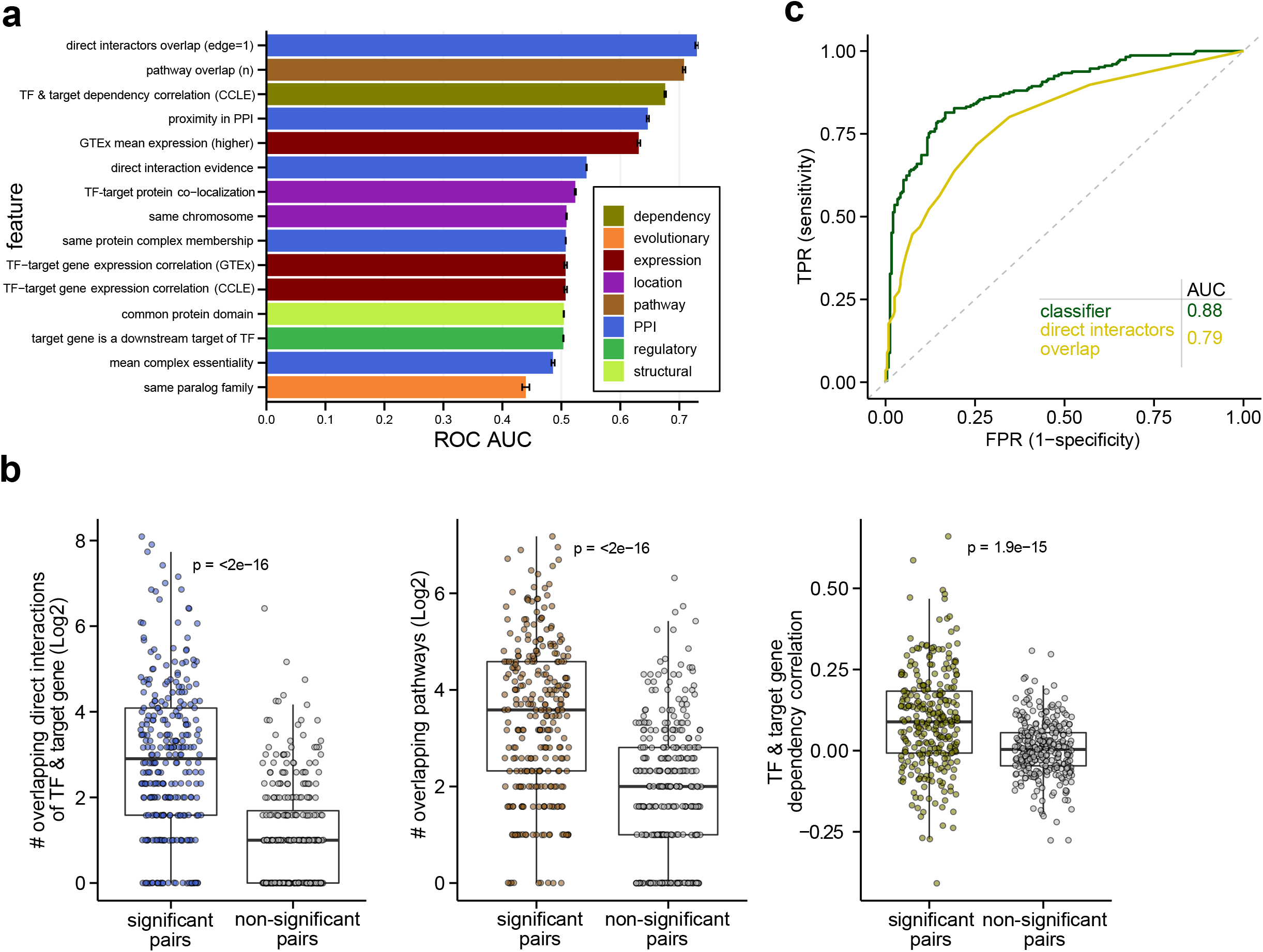
Features of TFa-target pairs. a) ROC AUC values of 15 different features in classifying TFa-target significant vs non-significant pairs. ROC AUC values were calculated by treating each feature as an independent classifier. b) Box plots illustrating the difference of top features from (a) between significant and non-significant pairs. “# of overlapping direct interactors”: overlap between the proteins that are directly connected to TF and its target gene in a protein-protein interaction network, “# overlapping pathways”: overlap of pathways in which TF and its target are involved, “TF & target gene dependency correlation”: dependency correlation of TF and its associated target gene across cell lines. c) AUC of a random forest classifier trained on 15 features from (a), compared to the top feature as the individual classifier.

Finally, we tested if the combination of the selected features can improve the ROC AUC of “significant pairs” classification compared to either feature alone. To this end, we trained a random forest classifier with all 15 features together. A random forest classifier can capture the interaction among different features and weigh the contribution of each feature in predicting a TFa-target significant relationship at an individual pair level. We found that our random forest model trained with an ensemble of features outperforms the top individual feature classifier (ROC AUC=0.79) with an ROC AUC of 0.88 on training data, which is 80% of 572 pairs (and ROC AUC of 0.92 on test data) (Fig 2c). From the classifier, we computed shapely additive explanation^39^ (SHAP) values for each pair to estimate the contribution of each feature to individual pairs (Supp Table 8). The SHAP value is a method for determining the contribution of each feature to the overall prediction of a model. This is achieved by evaluating the model’s performance with and without each feature and averaging the results across all possible permutations of feature ordering to ensure a fair comparison. The average contribution values of each feature across all pairs suggests that top features were consistent with individual classifier analysis (Supp Fig 2c).

Overall, we collected a set of features that capture the potential underlying biological features of TFa-target “significant pairs”. Our analysis revealed that the association between the TF and its predicted target is supported by involvement in shared molecular interactions and biological pathways.

### Cancer type specific TFa-target associations

Recent analyses^40^ focusing on identifying synthetic lethal pairs using mutational and dependency data emphasized the need to perform pan-cancer and individual cancer type focused analysis to identify associations that are missed by either analysis alone or to assign associations to specific cancer types. For example, in our analysis, we found NFE2L2 (also called NRF2) to be a strong self-dependent TF in pan-cancer analysis. An in-depth look into within cancer-types revealed the strongest correlation of NRF2 TF activity and its dependency in liver carcinoma (LIHC) and non-small cell lung carcinoma (NSCLC) compared to any other cancer type (Supp Fig 3a). In addition, certain TFs are known to involve cell identity of lineages and tumor specific expression programs to drive tumor maintenance and progression ^41,42^. Therefore, we reasoned that besides pan-cancer analysis of gene dependencies associated with TF activities, cancer type specific analyses are necessary. To this end, we performed similar linear modeling analyses of gene dependencies and TF activities in a cancer-type specific manner. We considered 26 cancer types with dependency and expression data for at least 10 cell lines in Project Achilles and prioritized TFs for each cancer type based on their expression in that cancer type (Supp Table 9, see methods).

In total, we identified 1,466 unique TFa-target pairs with a negative effect size (I.e., increased dependency of target with increased TFa) (FDR < 5%) across different cancer types comprised of 143 TFs and 715 target gene dependencies, which also included TF dependencies. The number of TFa-target pairs identified across different cancer types was highly variable, spanning from a single association in neuroblastoma (PATZ1-PFDN1) to 245 associations in breast cancer (Fig 3a, Supp Table 10). We further investigated the variability in the total number of TFa-target pairs identified across different cancer types and observed that the total number of identified pairs is correlated with the number of cell lines for which data is available (Supp Fig 3b). This suggests that the low number of cell lines for some cancer types could lead to an underpowered analysis, thus preventing the identification of statistically significant candidates. Among the cancer type specific TFa-target pairs, strongest associations include cases such as self-dependent TFs e.g., NFE2L2 in NSCLC and LIHC, STAT3 in lymphoma, and PAX5 in myeloma (Supp Table 10). Other strong interactions included GATA1-ZFPM1^43^ in leukemia and ZEB1-ITGAV in bladder carcinoma (Supp Table 10). Next, we compared if any of the TFa-target pairs span two or more cancer types and found that, on average 95% of the pairs identified in each cancer type are unique (Fig 3b). Only ∼3% (Supp Fig 3c) of pairs identified within individual cancers overlapped with TFa-target pairs from the pan-cancer analysis. This emphasizes the importance of performing cancer type analyses to expand the TFa-target space and identify context-specific TFa-target pairs.

**Figure 3:**
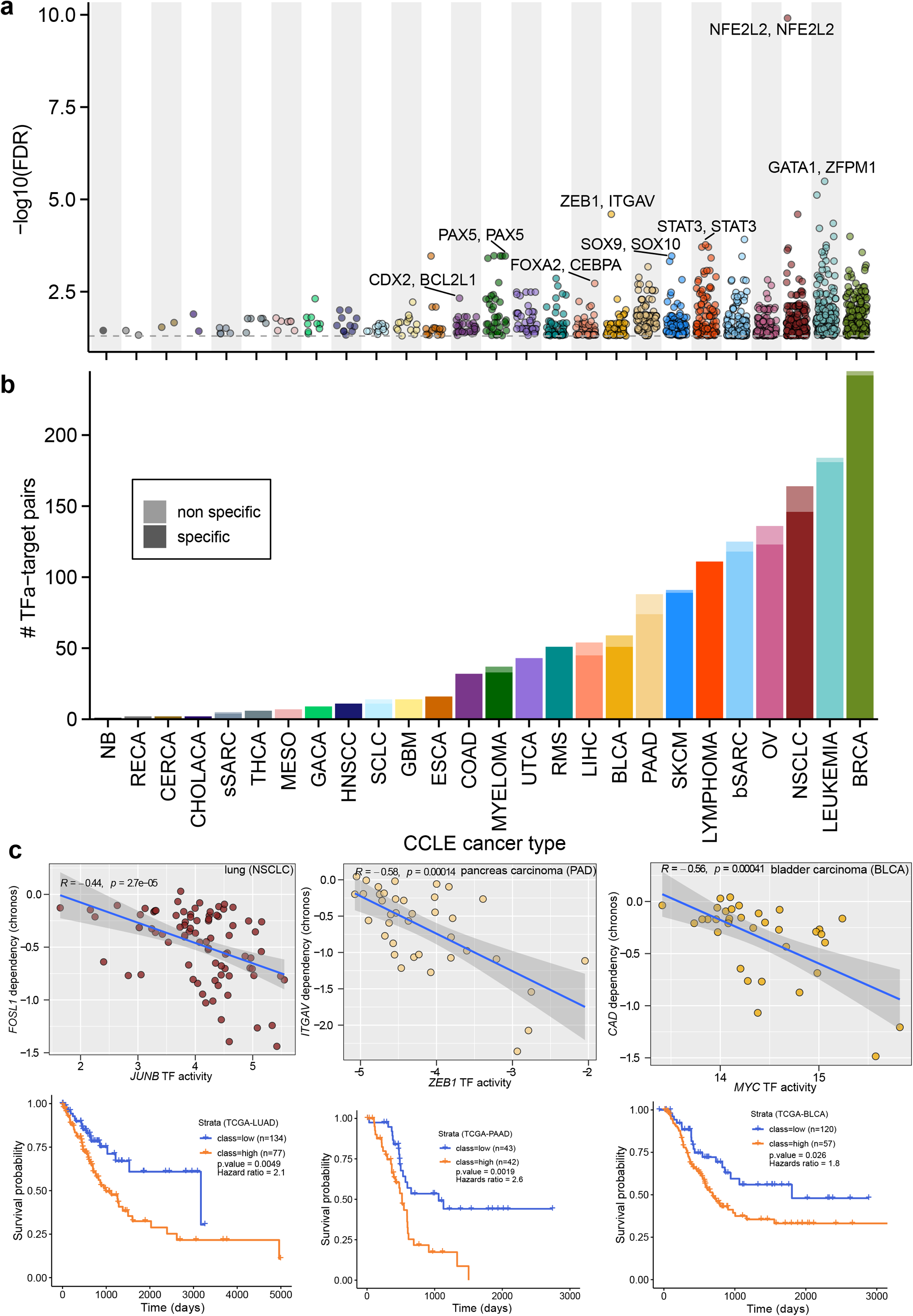
TFa-target pairs identified in individual cancer types. a) Manhattan plot showing the number of significant TFa-target pairs with negative effect size identified from each individual cancer type with the linear model-based framework (FDR < 5%). Top pairs from different cancer types are labeled. b) Bar plot showing the number of specific (dark color) and non-specific (light color) associations within each cancer type. c) Specific examples TFa-target pairs (JUNB-FOSL1, ZEB1-ITGAV, MYC-CAD) identified from different cancer types and their associated Kaplan-Meier plots of the same cancer type from TCGA.

To test the clinical relevance of cancer type specific pairs, we performed survival analysis using TCGA clinical and expression data as described in Supp Fig 1b. In total, 82.6% of the pairs are associated with poor survival in at least one TCGA cancer type when TF activity and its associated target gene expression both were high (Supp Fig 3d). Interestingly, only a small fraction of TFa-target pairs from cancer type specific analysis showed poor survival in matching TCGA cancer type (Supp Fig 3e). Among those, JUNB-FOSL1 pair was identified from lung cancer cell lines and this pair also showed poor prognosis when JUNB TF activity and FOSL1 expression was high in TCGA lung tumors (TCGA-LUAD). Similarly, ZEB1-ITGAV pair was identified in pancreatic carcinoma cell lines and showed poor prognosis in TCGA pancreatic adenocarcinoma (TCGA-PAAD) and MYC-CAD was identified in bladder carcinoma cell lines and showed poor prognosis in TCGA bladder carcinoma (TCGA-BLCA) (Fig 3c, Supp Table 11).

In total, the pan-cancer analyses across solid and hematological cancer cell lines and the cancer type specific analyses, resulted in 1,770 candidate TFa-target pairs comprising 151 TFs and 786 targets. Among these, 22 (∼1.24%) pairs are self-dependent TFs, 186 (∼10.5%) pairs include TFs as targets (i.e., TF activity is associated with other TF dependencies) and 1,562 (88.24%) pairs include non-TF genes as targets. To identify targets that could be suitable for pharmacological inhibition, we annotated target genes in our analysis with their target tractability information based on published literature. Target tractability indicates the likelihood of identifying a modulator that interacts with the given protein of interest^44^. For each of the 786 target genes, we extracted information about drug modality, inhibitor or antibody and tractability from literature^29,44^. Among the 786 targets from our analysis, currently 57 (∼7.2%, Supp Fig 3f) have inhibitors or antibodies targeting them. These include EGFR (inhibitors such as afatinib, erlotinib), JAK2 (ruxolitinib) and PIK3CA (taselisib). Next, we tested if our associations, based on genetic depletion and TFa, were also captured in pharmacological screens. To this end, we collected inhibitor response data in cell lines from the large-scale PRISM repurposing screen^45^ and correlated inhibitor responses with TF activities. EGFR inhibitor responses from PRISM database correlated positively with KLF5, FOXA1, GATA3 TF activities showing increased inhibitor sensitivity with increased TFa. This is in line with our TFa and dependency linear modeling analysis and suggests that the associations identified using genetic dependency data are recapitulated by pharmacological inhibition data. Similarly, we found PIK3CA to be associated with FOXA1 and TFAP2C TF activities across solid cancer cell lines (Supp Table 2). In agreement with these results, the pharmacological targeting of PIK3CA with taselisib, also showed a positive correlation with FOXA1 and TFAP2C TF activities (Supp Fig 3g, spearman correlation p.value < 0.05). For the remaining 729 targets, for which no inhibitor is available, we assigned previously defined target tractability buckets^29,44^. These tractability buckets ranged from 1 to 10, with 1 indicating the highest and 10 being the lowest tractability. 202 targets genes that are in this group have existing evidence to support their tractability but do not currently have any inhibitors available. The remaining targets (527/729, ∼72.3%) were assigned to buckets 8-10 (Supp Fig 3f). Targets in this group have little to no existing evidence to support their tractability. Across the identified 786 target genes, we estimated that more than 27% presented characteristics for pharmacological molecule development, recommending them for further investigation.

In summary, we identified a range of TFa-target pairs that are specific to different tumor types by performing cancer-type focused linear modeling analysis of TF activity and dependency analysis. A small fraction of targets associated with different TF activities currently have an existing inhibitor. Using the large-scale inhibitors response data from PRISM database, we have shown that genetic dependency associations with TF activities can be translated to inhibitor responses.

### *TEAD1* activity associated gene dependencies

The TEA domain (TEAD) containing family of TFs is comprised of four paralogues (TEAD1-4) in humans, which together with their established cofactors YAP and TAZ (WWTR1) are implicated in tumor progression, metastasis, and therapy resistance in a broad range of cancer types^34,46–55^. Consistently, in our pan-cancer analysis we identified TEAD1 to be associated with a large number of gene dependencies (Fig 1c). In total, we identified 27 targets whose increased dependency is correlated with increased TEAD1 activity (Fig 4a). Among these, TEAD1 itself was identified as a vulnerability, suggesting TEAD1 as a self-dependent TF. Other targets included previously established transcriptional coactivators of TEAD such as the paralogues WWTR1/TAZ, YAP^56^, and genes described in the context of hippo pathway e.g., PTK2 or ITGAV^57^ (Supp Table 12). To understand the relevance of the identified target genes, we performed pathway enrichment with the 26 genes (excluding TEAD1) and found the terms “Hippo pathway”, “focal adhesion” and “TGF-beta signaling pathways” as significantly enriched (Supp Fig 4a). These observations are in line with previous evidence of a cross talk between the TGF-beta and Hippo pathways^58,59^.

**Figure 4:**
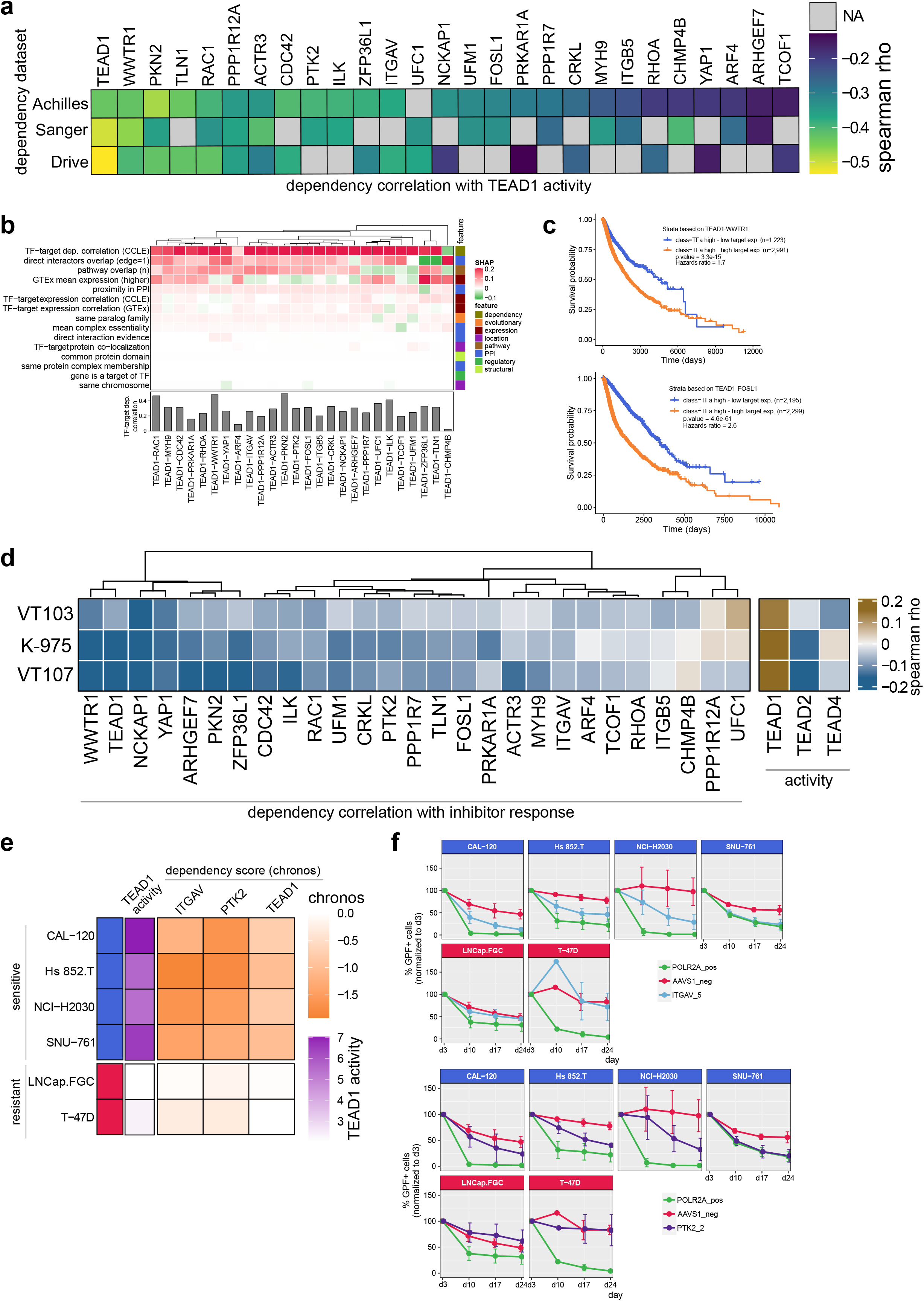
TEAD1 TFa associated gene dependencies & validation. a) Heatmap showing the correlation of 27 target gene dependencies with TEAD1 TFa (spearman correlation, 95% confidence intervals). b) Heatmap of Shapley Additive Explanation (SHAP) values of all features of TEAD1 TFa associated targets extracted from random forest classifier explained in Fig 3c. X-axis shows all 26 genes (excluding TEAD1 dependency). c) Example Kaplan-Meier plots showing the survival difference between low vs high WWTR1 or FOSL1 expressing subgroups with high TEAD1 TFa. Subgroups were defined same as Fig 1g. d) Heatmap showing the correlation of TEAD1 TFa target gene dependencies with inhibitor responses and TEAD paralogue activities (spearman correlation, 95% confidence intervals). VT103 – TEAD1 specific inhibitor, VT107 – pan-TEAD inhibitor, K-975 – YAP/TAZ interaction inhibitor. e) Predicted sensitive and resistant cell lines selected for CRISPR/Cas9 depletion assays. TEAD1 activity and PTK2, ITGAV and TEAD1 dependency scores are shown. f) CRISPR/Cas9 depletion time course showing relative growth defects introduced by sgRNAs against PTK2 (purple), ITGAV (blue), the negative control AAVSI (red), or the positive control sgRNA POLRA (green). Guide containing GFP+ cell numbers of weekly measurements over 24 days were normalized to the GFP+ cell numbers of day 3 upon transduction with the guides. Mean and standard deviation of three biological replicates are shown. For T-47D resistant cell line, two biological replicates were presented and with one measurement on day 10.

Next, we investigated the Shapley additive explanation (SHAP) values that were annotated based on the random forest classifier described above to assess the 15 features contributing to these 26 target gene dependencies associated with TEAD1 activity. For the majority of the target genes, the highest ranking feature was the strong positive correlation between the cell lines’ dependency on the target as well as the dependency on TEAD1 (Fig 4b). This observation suggests that tumors that are dependent on TEAD1 are also likely dependent on either one or a combination of these target genes. Next, we calculated the correlation of TEAD1 dependency with dependencies of all genes across the 757 cell lines. Among the top 50 genes with positive correlation, we observed 12 of our TEAD1 activity associated target genes were overlapping (Supp Fig 4b), suggesting that additional vulnerabilities were revealed from our analysis based on TEAD1 activity.

Next, we tested the impact of the 26 target genes and TEAD1 activity in TCGA patient tumors. Of the 26 TEAD1 target genes, 24 showed poor prognosis in at least one cancer type when expressed together with high TEAD1 TFa (Fig 4c, Supp Fig 4c, Supp Table 3). In addition to the analysis of the individual expression of the 26 target genes, we combined their expression using single sample gene set enrichment analysis to compute a score for each tumor sample in TCGA. We tested the correlation between the expression-score of the targets and the activity scores of three of the TEAD members with high confidence regulon annotation across the patient samples. We found that the expression score of the 26 genes is positively correlated specifically with TEAD1 activity as opposed to TEAD2 or TEAD4 activity, suggesting the specificity of regulons in defining TEAD1 activity and its association with predicted target genes (Supp Fig 4d). We further tested the specificity of the 26 target-associations across the TEAD family members in cell line data. We computed expression-score similarly for all cell lines for which dependency data is available and divided cell lines into three equal sized subgroups (low, medium, high) and compared TEAD paralog dependencies across these three subgroups of cell lines. This analysis revealed that cell line subgroup with high expression-score also showed strong TEAD1 dependency compared to other TEAD paralog dependencies (Supp Fig 4e), confirming the specificity of these target genes to TEAD1. These results confirmed that the associations of the TEAD1 activity and the 26 predicted target genes are recapitulated across multiple data domains and highlighted a TEAD1-specific association among the different TEAD paralog family members.

TEAD family members are among the few TFs for which pre-clinical inhibitors are currently available. We hypothesized that pharmaceutical targeting of the TF itself will recapitulate the results of our analysis and that increased responses (i.e., anti-proliferative effect) of these inhibitors will not only correlate with TEAD1 activity but also correlate with increased target gene dependency. To this end, we profiled the pharmacological impact of three inhibitors in 757 solid cancer cell lines using PRISM inhibitor screening approach^45,60^. For this experiment, we used VT103 (a TEAD1 specific inhibitor), VT107 (a pan-TEAD inhibitor) and K-975 (YAP/TAZ interaction inhibitor)^61,62^ (Supp Table 13). Although the anti-proliferative effect (measured as activity area, i.e., the area above the fitted dose response curve) elicited by these inhibitors was not very strong across cell lines, we still identified a positive correlation of inhibitor response and TEAD1 activity (i.e., increased inhibitor response with increased TEAD1 activity). Similarly, a negative correlation of inhibitor response and target gene dependency was observed (i.e., increased inhibitor response with increased target gene dependency) (Fig 4d). Taken together, the target gene dependencies that are associated with TEAD1 activity show similar correlation with pharmacological inhibition of TEAD.

Finally, through CRISPR-Cas9 depletion assays, we experimentally tested two of our TEAD1 associations, ITGAV and PTK2. To this end, we selected six cell lines and classified them as sensitive (Cal120, Hs852T, NCI-H2030, SNU761) and resistant (LNCap and T-47D) based on TEAD1 activity and ITGAV, PTK2 dependency (Fig 4e). We transduced the cell lines with Cas9 and gRNAs targeting ITGAV, PTK2 and TEAD1 plus a GFP reporter. The gRNAs induced a clear loss of target gene expression at day 7 after transduction (Supp Fig 5a). The relative proliferation of GFP positive cells within total cell population was monitored from day 3 until day 24 by flow cytometry. For both target genes ITGAV and PTK2, both sgRNAs we tested caused a strong reduction cell proliferation in cell lines classified as sensitive, while in resistant cell lines this effect was not observed (Fig 4f, Supp Fig 5b, Supp Table 14). In addition, we also observed a similar reduction in cell proliferation with sgRNAs targeting TEAD1, thus also validating the self-dependency of TEAD1 in cell lines classified as sensitive (Supp Fig 5b). These results confirm that the cell lines we classified based on TEAD1 activity as a biomarker are sensitive to loss of PTK2 or ITGAV, two of our predicted target genes associated with TEAD1 activity (Fig 4f). Finally, we tested if the results obtained via genetic loss-of-function mediated by CRISPR deletion can be recapitulated by pharmacological inhibition of TEAD. To this end, we treated the sensitive cell lines with the pan-TEAD inhibitor VT-107^61^ (Tang et al.) and the ITGAV antibody Intetumumab and quantified the changes in the expression level of the canonical TEAD downstream target CTGF by RT-qPCR. As expected, we observed a dose dependent reduction of CTGF expression with the inhibitor and antibody in most cell lines (Supp Fig 5c). These results validate the cell lines selection based on their TEAD activity and recapitulate the genetic inhibition observations with pharmacological inhibition of ITGAV and TEAD (Supp Fig 5d).

In summary, we identified a set of 26 target gene dependency associations for TEAD1 activity, two of which we validated experimentally through CRISPR-Cas9 depletion assaysand pharmacological inhibition. Further, we confirmed that target gene dependencies associated with TEAD1 activity are also similarly associated with TEAD inhibitor responses.

## Discussion

Efficient therapies in oncology rely on the identification of specific and robust biomarkers and targets. Several computational methods have used pan-cancer genetic perturbation and matched genomic alterations data (e.g., mutations, copy number alterations, microsatellite instability) to establish potential biomarker-target pairs in the past^27,29,30,40^. Recently, we and others have shown the power of quantitative omics data (e.g., gene expression) combined with genome-wide perturbation data or exclusively gene expression data together with patient survival data in identifying potential gene-gene interactions leading to synthetic lethality (SL)^37,63^. SL interactions indicate cell death through the co-inactivation of both genes in a pair whereas the inactivation of either gene alone does not affect viability ^64–66^. However, a focused analysis of TF activities (TFa), key components in regulating tumor-specific transcriptional programs, and their associated genetic vulnerabilities is lacking. In the case of TFs, in addition to genomic alterations, their activity is impacted by complex regulatory mechanisms involving multiple proteins and genesas we **l** as epigenetic and post-translational modifications. In this work, we derived TF activities using their annotated regulons expression^32^ and used them as potential biomarkers in our linear model framework to identify associated genetic dependencies (targets). Our approach, using TFa complements prev ious biomarker assessments through discrete genomic alterations. Our analysis expands beyond classical SL, by identifying targets whose inhibition will lead to a lethal phenotype in the presence of a hyperactive TF, previously described as synthetic dosage lethality^64,67,68^. We propose that these interactions are highly relevant in the current medical context, where direct pharmacological targeting of oncogenic TFs is still highly challenging, and identification of novel TF vulnerabilities can lead to a strong therapeutic impact.

We identified a total of 1,770 unique TFa-target candidate pairs, for which increased target dependency is associated with increased TFa. We hypothesize that these candidate pairs represent a broader impact for further validations. For example, ZEB1-ITGAV pair was identified in bladder carcinoma, non-small cell lung carcinoma (NSCLC) and pancreatic carcinoma (PAAD) cell lines. Though direct interaction of these two proteins was not previously reported, both are involved in epithelial-to-mesenchymal transition (EMT), with ZEB1 as a key TF in EMT initiation^69,70^, while ITGAV is EMT associated cell surface marker. Another candidate pair RUNX2-JUN was exclusively identified in glioblastoma cell lines. However, we observed survival impact of this pair not only in glioblastoma but also in low-grade glioma and bladder carcinoma (Supp Table 11). The proteins encoded by these two genes were reported to physically interact and regulate the expression of downstream genes and pathways^71,72^. Though the candidate pairs reported in our study require additional experimental validations, as exemplified they hold great promise as potential biomarker-target pairs.

Across our different candidate pairs, TEAD1 emerged at the center of multiple interactions. TEAD1 and its paralog family members (TEAD1-4), and their co-factors YAP and TAZ mediate the transcriptional output downstream of the Hippo signaling pathway, which is involved in regulating cell proliferation and migration. With a focus on TEAD1 activity as a biomarker and its set of 26 predicted targets, we designed additional experiments to understand the observed interactions. TEADs are among the small number of TFs for which pharmacological inhibitors are available, both specifically targeting TEAD1 (VT103), pan-TEAD (VT107) and YAP/TAZ interaction inhibitor (K975)^61,62^. Previous studies integrating perturbation and drug sensitivity measurements across cell lines identified drug mechanism of action and drug’s secondary targets^73,74^. Similar to these observations, our results show that TEAD1 target dependencies are correlated with TEAD inhibitor sensitivity, indicating the effect of direct pharmacological targeting of the TEAD TFs. In addition, with single gene depletion assays, we showed that cell lines with high TEAD1 activity are vulnerable to ITGAV or PTK2 genetic inhibition. This is in accordance with recently published literature showing a feedback regulatory loop between TAZ/ITGAV and YAP/TAZ in Hippo pathway activation^57^ and focal adhesion kinases (PTK2) modulating YAP/TAZ activity^75^. With growing evidence on the role of Hippo signaling pathway in cancers, our TEAD1-specific analysis provides an example of relevant new clinical targets and drug-repurposing strategies for further therapeutic exploration.

While we made efforts to increase the robustness of our predictions based on integration across independent genetic depletion screens, stringent statistical approaches, and assessment of clinical relevance across patient cohorts, our approach is subject to several limitations. First, while cell lines have proven to be a useful tool in cancer research, they do have limitations related to their representation of primary tumors and the translation of results to the *in vivo* context. While we systematically use TCGA patient cohorts for a first assessment of the predicted interaction in a clinically relevant context, the specificity of each one of our selected pairs in defining patient populations would need validation across independent cohorts that are currently not covered in this study. Second, the power of our analysis is impacted by the number of cell lines. In our cancer-type focused analyses, the number of cell lines varies between 13 (mesothelioma) to 84 (NSCLC). Specifically, we expect cancer types represented by a small number of cell lines to fall short of providing the expected statistical power for the comprehensive identification of candidate TFa-target pairs. Third, the underlying molecular mechanisms of the predicted associations need in-depth experimental work beyond the purpose of this study. Further improvements of our methodology could integrate protein abundance quantification in addition to gene expression, as well as expand the current one TF to one target pairs to interactions between multiple TFs and their respective targets.

Ongoing developments in experimental approaches such as combinatorial screens ^76^ or drug anchored screens are providing the required tools to map complex genetic interactions in cells. The method and validations we provided in this work is a step toward understanding such interactions and potentially identifying new targets and biomarkers involving TFs. The targets we identified for each TF can serve as the basis for designing targeted libraries for above mentioned experimental strategies and highlight the power of TFa to stratify the tumors and harness their underlying vulnerabilities for future targeted therapeutics.

## Methods

### Transcription factor activity (TFa) estimation in cell lines and TCGA tumors

We estimated a given transcription factor activity (TFa) using the combined expression of its annotated target genes, called regulons. Regulons for each TF were extracted from the dorothea R/Bioconductor package^32^, which includes a collection of annotated TF targets from different sources^32^. We then computed the activity score for each TF using the Viper R/Bioconductor package. The TF activity is defined as the normalized enrichment score (NES) calcuated by the Viper algorithm^77^.

### Pan-cancer identification of TFa associated gene dependencies (TFa-target pairs)

We downloaded a list of human transcription factors from reference^78^ and selected transcription factors (TFs) that have regulon annotations in Dorothea R/Bioconductor package. We removed TFs which do not show the dependency (dependency score <=-0.8) in at least three cell lines in two out of three dependency databases (Project Achilles^27^, Project Sanger^29^, Project Drive^30^) and TFs which show dependency in >=95% of cell lines included in Project Achilles (Supp Fig 1a). We selected TFs in this way to remove “never essential” and “pan-essential” TFs from our analysis and keep those TFs that show a potential context dependency. We used the resulting 92 TFs to identify their activity-associated gene dependencies (Supp Fig 1a). To identify the gene dependencies associated with TFa, we implemented a linear regression model using the formula “*gene dependency ∼ TFa + Cancer type*”. Cancer type of each cell line was included in order to account for the variation in dependency scores due to different tissue types. We implemented this linear model for each TF with all genes (including all TFs) as potential target genes. We then adjusted nominal p-values for multiple testing using the Benjamini-Hochberg method after excluding “pan and never essential” genes, keeping 11,665 dependencies as potential targets. This procedure has been repeated with three cancer dependency datasets Project Achilles, Sanger and Project Drive. We called TFa-target pair significant if the adjusted p.value was < 0.1 (FDR < 10%) and the TFa coefficient (effect size) was above or below 3 standard deviations. TFa-target pairs with negative effect size indicate increased dependency on a target gene with increased TFa and positive effect size indicates increased dependency on a target gene with decreased TFa. We considered TFa-target pairs with negative effect size for the remaining downstream analysis.

### Cancer type specific TFa-target pairs identification

To identify cancer-type specific TFa-target pairs, we selected those cancer types for which expression and dependency data are available for at least 10 cell lines in Project Achilles.^27^ We chose this CRISPR KO dataset due to the inclusion of a large number of cell lines. For each cancer type, to remove ‘pan-essential’ and ‘never essential’ TFs, we removed TFs which showed dependency (chronos score <= -0.8) in >95% of cell lines or did not show dependency in at least one cell line. Among the remaining TFs, we selected the TFs based on their expression. We computed the mean for each TF across all cell lines for a given cancer type and divided the expression of all TFs into three equal sized bins (tertiles). We then considered TFs falling into the second and third tertiles as the expressed TFs for that cancer type (Supp Table 9) and computed their activities. Similar to TF dependency, we removed gene dependencies that did not show dependency in at least one cell line and showed dependency in >95% of cell lines and kept the remaining gene dependencies as the potential target pool.

We implemented a linear model to identify TFa-target pairs in the same way as described above for the pan-cancer analysis, except in this analysis we did not use cancer type as a covariate. We then adjusted nominal p-values for multiple testing using Benjamini-Hochberg method and considered TFa-target pairs with FDR < %5 and TFa coefficient (effect size) above or below two standard deviation values to reduce the number of false positives. TFa-target pairs with negative effect size indicate increased dependency on a target gene with increased TFa and positive effect size indicates increased dependency on a target gene with decreased TFa. We considered TFa-target pairs with negative effect size for the remaining downstream analysis.

### TCGA cohort definitions and survival analysis

To assess the clinical relevance of identified TFa-target pairs with negative effect size from both pan-cancer and cancer type specific analysis, we used TCGA expression and clinical data to perform survival analysis as illustrated in Supp Fig 1b. For each TCGA cancer type, we computed the TF activity with the dorothea and viper R/bioconductor packages as described above. For a TFa-target pair, we divided each cohort into three equally sized subgroups (low, medium, high) based on TFa and target gene expression independently. We removed the tumors in the ‘low’ TFa subgroup and defined the tumors in the ‘medium’ and ‘high’ tertiles as the ‘high’ TFa subgroup. We then overlapped these high TFa subgroups of tumors with tumors in ‘low’ and ‘high’ target expression subgroups. Finally, we defined TFa high and low target expression (TFa high – low target expression) and TFa high and high target expression (TFa high – high target expression) subgroups and compared the survival difference between these two with a cox proportional hazards model. We performed this analysis on each TCGA cancer type and in a pan-cancer setting by combining all solid TCGA cancers together. For the pan-cancer analysis, we used cancer type as the covariate in the cox proportional hazards model. We used coxph function from survival R/biocondcutor package and survival curves shown in the figures were plotted with ggsurvplot from survminer R/bioconductor package. We considered a survival difference between compared subgroups significant, if both subgroups contain at least 10 tumors, the p-value was < 0.05 and the hazards ratio was > 1 for “TFa-high – high target expression” subgroup compared to “TFa-high – low target expression” subgroup.

### Ranking of TFa-target pairs features and random forest classifier implementation

To identify the potential features underlying TFa-target pairs with negative effect size, we defined a set of 15 features that potentially capture the relationship of two genes at different levels. All features and their sources are listed in Supp Table 7. These features contain different layers of information such as protein-protein interactions, expression level association (correlation of expression), evolutionary relatedness (meaning belonging to the same paralog families), and shared pathway membership. We prepared the dataset of TFa-target pairs that were significant and non-significant with negative effect size from solid pan-cancer analysis and labeled them as “significant pairs” or “non-significant pairs”. We removed self-dependent TFs (where TFa and target are same) and TFa-target pairs where target is also a TF. For this combined set of 572 pairs (272 significant pairs and 300 randomly selected non-significant pairs), we first computed the ROC AUC of each feature with the roc function from pROC R package to identify the features with top ranking. To identify the contribution of each feature to each significant pair, we trained a random forest model with an ensemble of 15 features and all 572 TFa-target significant and non-significant pairs. The random forest classifier model was trained using the randomForest R package using 80% of the whole dataset as training data and 3 repeat cross fold validation. We further computed the contribution of each feature to each pair as Shapley additive explanations (SHAP) values using the treeshap R package (Supp Table 8).

### GSEA/enrichment analysis

Gene set enrichment analysis (GSEA) analysis^79^ has been performed with the clusterProfiler R/Bioconductor package^80^ using gene sets from the molecular signatures database^81^ (MSigDB). The nominal p-values were adjusted using the Benjamini-Hochberg method and significant enrichments were defined as having an adjusted p.value < 0.05.

### Expression-score for TCGA tumors with 26 genes whose dependency is associated TEAD1

We used the gene set variation analysis (GSVA) R/bioconductor package to compute expression-score for all TCGA tumors using target genes associated with TEAD1 as a gene set. We first removed WWTR1/TAZ and YAP1 genes from 26 target genes of TEAD1 before computing expression-score for all TCGA tumors, since these two genes were also annotated as regulons of TEAD1 to avoid bias in the expression-score. This expression-score was correlated with TEAD1 activity across different TCGA cancer types.

### PRISM inhibitor screen

We performed a PRISM inhibitor screen using the TEAD1 specific inhibitor (VT103), the pan-TEAD inhibitor (VT107) and the YAP/TAZ interaction inhibitor (K-975) as described previously^45, 60^. These assays were performed at the Broad Institute; inhibitors were synthesized and analyzed for quality as described in literature^61,62^. Cell lines were grown in RPMI 10% FBS without phenol red for adherent lines and RPMI 20% FBS without phenol red for suspension lines. Parental cell lines were stably infected with a unique 24-nucleotide DNA barcode via lentiviral transduction and blasticidin selection. After selection, barcoded ce **l** lines were expanded and QCed (mycoplasma contamination test, a SNP test for confirming cell line identity, and barcode ID confirmation). Passing barcoded lines were then pooled(20-25 cell lines per pool) based on doubling time and frozen in assay-ready vials.

Test compounds were added to 384-well plates and run at 8 pt. dose with 3-fold dilutions in triplicate with a top dose of 10uM. These assay-ready plates were thenseededwith the thawed cell line pools. Adherent cell pools were plated at 1250 cells per well, while suspension and mixed adherent/suspension pools were plated at 2000 cells per well. Treated cells were incubated for 5 days, then lysed. Lysate plates were collapsed together prior to barcode amplification and detection.

Each cell line’s unique barcode is located at the end of the blasticidin resistance gene and gets expressed as mRNA. These mRNAs were captured using magnetic particles that recognize polyA sequences. Captured mRNA was reverse transcribed into cDNA and then the sequence containing the unique PRISM barcode was amplified using PCR. Finally, Luminex beads that recognize the specific barcode sequences in the cell set were hybridized to the PCR products and detected using a Luminex scanner which reports signal as a median fluorescent intensity (MFI).

### PRISM inhibitor screen data processing

For each plate, we first normalized the logMFI (log2 mean fluorescence intensity) of the DMSO wells to their median logMFI. Each detection well contained 10 control barcodes in increasing abundances as spike-in controls. A monotonic smooth p-spline was fit for each control barcode detection well to normalize the abundance of each barcode to the corresponding value in the plate-wise median DMSO profiles. Next, all the logMFI values in the well were transformed through the inferred spline function to correct for amplification and detection artifacts. Next, the separability between negative and positive control treatments was assessed. In particular, we used the error rate of the optimum simple threshold classifier between the control samples for each cell line and plate combination. Error rate is a measure of overlap of the two control sets and is defined as “*Error=(FP+FN)/n*” where FP is false positives, FN is false negatives, and n is the total number of controls. A threshold was set between the distributions of positive and negative control logMFI values (with everything below the threshold said to be positive and above said to be negative) such that this value is minimized. Additionally, we also filtered based on the dynamic range of each cell line. Dynamic range was defined as “*DR= μ*_*-*_ *- μ*_*+*_” where μ+/− stood for the median of the normalized logMFI values in positive/negative control samples. We filtered out cell lines with error rate above 0.05 and a dynamic range less than 1.74 from the downstream analysis. Additionally, any cell line that has less than 2 passing replicates was also omitted for the sake of reproducibility. Finally, we computed viability by normalizing with respect to the median negative control for each plate. Log-fold-change viabilities were computed as “*log-viability = log2(x)-log2(μ*_*-*_)” where *log2(x)* is the corrected logMFI value in the treatment and *log2(μ*−) is the median corrected logMFI in the negative control wells in the same plate. Log-viability scores were corrected for batch effects coming from pools and culture conditions using the ComBat algorithm as described previously^82^. We fit a robust four-parameter logistic curve to the response of each cell line to the compound: *f(x) = b+((a-b)/1+e*^*slog(x/EC50)*^) with the following restrictions 1) We require that the upper asymptote of the curve be between 0.99 and 2) We require that the lower asymptote of the curve be between 0 and 1.01 3) We no longer enforce decreasing curves 4) We initialize the curve fitting algorithm to guess an upper asymptote of 1 and a lower asymptote of 0.5 5) When the standard curve fit fails, we now report the robust fits provided by the dr4pl R-package and computed AUC (area under the curve) and IC50 values for each dose-response curve. Finally, the replicates were collapsed to a treatment level profile by computing the median score for each cell line. Finally, the activity area for each inhibitor across cell lines was correlated with target gene dependency and TEAD paralogue activities using corr function in R.

### Cell lines

Cell lines were obtained from cell banks as indicated in the resource table. NCI -H2030, SNU761, LNCap.FGC and T47D were cultured in RPMI 1640 ATCC formulation supplemented with 10% FCS. The media of T47D was additionally supplemented with 0.2 U/ml bovine insulin. Hs852.T and Cal-120 were cultured in DMEM supplemented with 10% FCS. LentiX were propagated in DMEM supplemented with Tet-approved 10% FCS. All cell lines were cultivated at 37°C in a humidified atmosphere with 5% CO_2_.

### Generation of Cas9 expressing cell lines

The codon-optimized cDNA sequence of Cas9 was synthesized and cloned into backbones by GenScript Biotech Corporation ^83^. The Cas9 plasmid was packed into lentiviral particles using the LentiX cell line and LentiX VSV-G single shots according to manufacturer’s protocol. After 48h the lentiviral particle-containing supernatants were harvested, filtered through a 0.45 μm SFCA filter and stored at -80°C until further usage. NCI-H2030, SNU761, LNCap.FGC, T47D, Hs852.T, and Cal-120 were transduced with Cas9-containing lentivirus and selected 3 days later with puromycin.

### CRISPR/Cas9 depletion assays

All CRISPR/Cas9 depletion assays were conducted as previously described previously^84^. Briefly, gRNAs were cloned into the lentiviral vector Lenti_gRNA GFP(LRG)_2.1T by GenScript Biotech. Then lentiviral particles were produced in LentiX cells using LentiX VSV-G shots to package the guide containing lentiviral vectors, according to manufacturer’s protocol. Viral supernatants were harvested after 48h, filtered through a 0.45 μm SFCA filter, aliquoted and stored at -80°C until further usage. For depletion assays, relevant cell lines stably expressing Cas9 were seeded at approximately 50–60% in a 12-well plate and transduced with viral particles to achieve a transduction efficiency of 10-90%. Cells were then cultured for 24 days and were analyzed for GFP positive cells on the BD FACSCanto II flow cytometer on a weekly basis starting at day 3 after transduction.

### Fluorescence activated cell sorting (FACS)

NCI-H2030 cells were seeded in a 6-well plate and transduced with guide-containing lentiviral particles targeting TEAD1 and PTK2 with a transduction efficiency of 50-90%. Seven days upon transfection and expansion to T75 tissue-culture flask, cells were detached with Accumax and sorted with SONY Sorter SH800Z to gain 0,5-2*10^5 cells of a > 95% pure GFP+ cell population for respective guides. Cells were pelleted by centrifugation at 300x g for 5 min, washed once with PBS and lysed in RIPA lysis buffer plus complete and 0.5% dodecyl-β-D-maltoside.

### Inhibitors titrations and CTGF expression by qPCR

For the treatment with inhibitors, 10000 cells/well were seeded in a 96 well plate (96 well, white, clear F-Bottom, Lid, tissue-culture treated plates, sterile, PerkinElmer PNr.: 6005181). The day after seeding the cells, compounds were added using a digital dispenser (D300 Digital Dispenser, HP) in the case of the sma**l** molecule inhibitors, or added manually in the case of Intetumumab and respective IgG control. 48h after the beginning of inhibitors treatment, cells were harvested and processed using the FastLane Cell RT-PCR_QuantiTect Multiplex RT-PCR Kit (Qiagen #216513). For the amplification of CTGF and b2-Microglobulin we used TaqMan probes (B2M VIC, Cat. Nr. Hs00187842_m1, and CTGF FAM, Cat. Nr. Hs00170014_m1).

### Protein quantification by WES

Protein content of lysates was quantified with BCA assay according to manufacturer’s recommendations. Individual protein KO was determined using the automated Western Blot System WES from Proteintech. Therefore, 0.2 - 0.4 μg/μl lysate was loaded onto a 12-230 kDa 25-Capillary Cartridge and was incubated for 60 min with primary antibodies against TEAD-1 (1:50), PTK2 (1:200), ITGAV (1:200) and GAPDH (1:2000). After 30 minutes incubation with the anti-rabbit secondary reagent, signals were developed using chemiluminescence.

## Data availability

The list of human transcription factors (v1.0.1) were downloaded from http://humantfs.ccbr.utoronto.ca/download.php in August2021. Annotated transcription factor regulons were used from dorothea R/bioconductor package ^32^ (v1.0.1). The dependency data of Project Achilles, Project Sanger, Project Drive and expression data of CCLE cell lines was downloaded from https://depmap.org/ fromrelease 2021 Q4. The cancer genome atlas (TCGA) data was collected from GDC portal https://portal.gdc.cancer.gov/ and the expression data was processed as previously described^85^. Pathway gene sets were downloaded from molecular signatures database^81^ (https://www.gsea-msigdb.org/gsea/msigdb/) (v7.5.1). Protein-protein interactions were downloaded from the biogrid database (https://thebiogrid.org/) (v4.4.204). Target tractability data is collected from publication (https://www.nature.com/articles/s41586-019-1103-9). PRISM inhibitor screen performed as part of this study are provided for public access in Supp Table 13. All reagent sources used in CRISPR-Cas9 depletion assays are provided in Supp Table 15.

### Code availability

All analyses were performed in R statistical programming language (v4.0.2)

- Tidyverse (v1.3.0) - https://github.com/tidyverse/tidyverse
- Dorothea (1.0.1) - https://bioconductor.org/packages/release/data/experiment/html/dorothea.html
- Survival (v3.2-3) - https://cran.r-project.org/web/packages/survival/index.html
- Survminer (v0.4.8) - https://cran.r-project.org/web/packages/survminer/index.html
- pROC (v1.16.2) - https://cran.r-project.org/web/packages/pROC/index.html
- randomForest (v4.6-14) - https://cran.r-project.org/web/packages/randomForest/index.html
- Treeshap (v0.1.1) - https://github.com/ModelOriented/treeshap
- GSVA (v1.36.2) - https://bioconductor.org/packages/release/bioc/html/GSVA.html
- ComplexHeatmap (v2.4.3) - https://bioconductor.org/packages/release/bioc/html/ComplexHeatmap.html
- Code written for analysis in this paper – Code will be made open source upon the publication

## Figure Legends

**Supp Figure 1.**
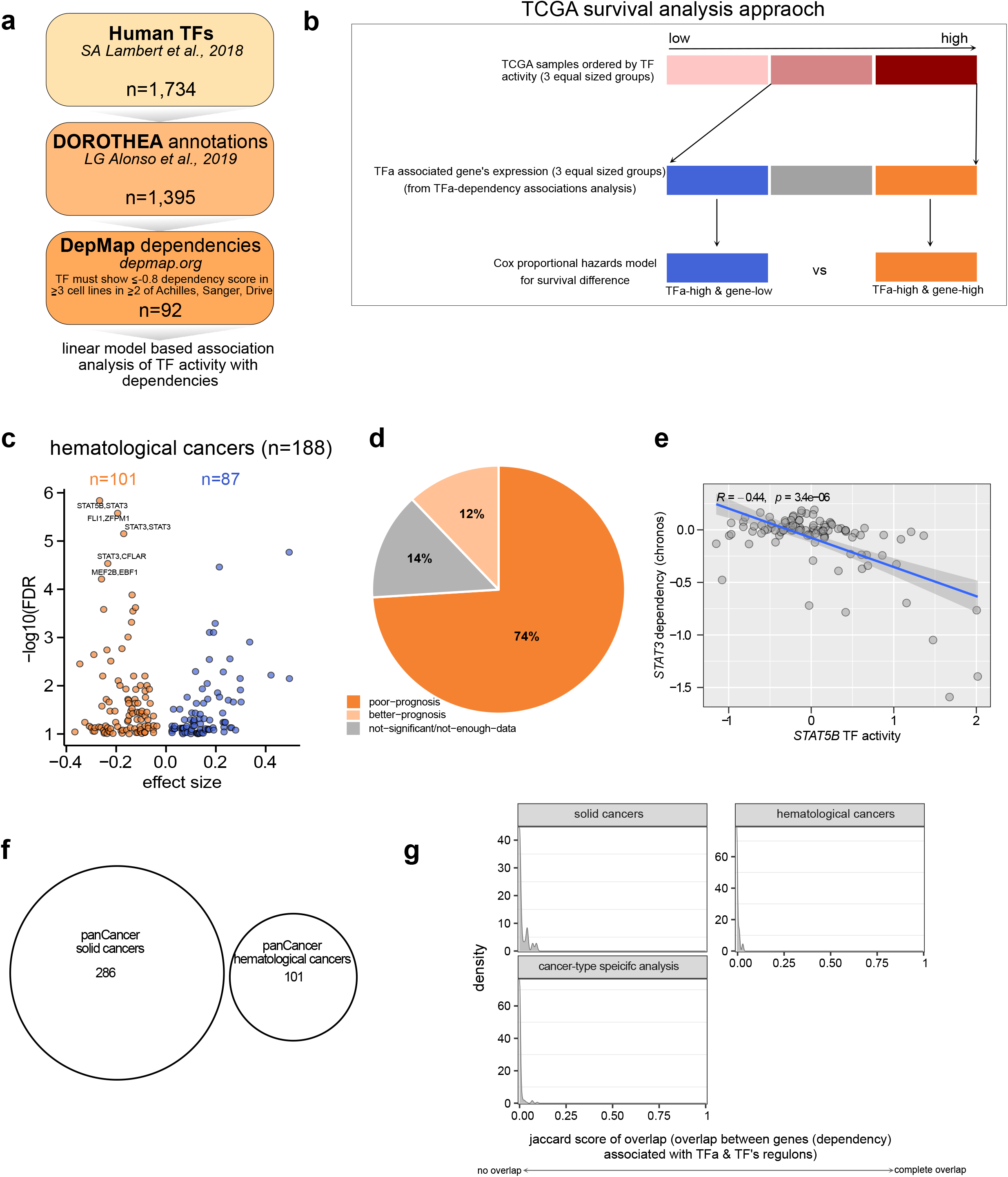
a) Flow chart illustrating the prioritization of transcription factors for linear model based analysis of 757 solid cancer cell lines. b) Illustration showing the approach employed for TCGA survival analysis for each TFa-target pair. This analysis was performed on all TCGA solid tumors together as pan-cancer setting and on each individual TCGA cancer type. c) Volcano plot showing the total number of TFa-target pairs identified from linear model based framework in hematological cancer cell lines. d) Pie chart indicating the fraction of TFa-target pairs from (c) and their association with survival of TCGA cancer types (same as Fig 1e). e) Scatter plot showing the correlation of TFa and dependency of the top TFa-target pair, STAT5B-STAT3, from hematological cell lines (spearman correlation, 95% confidence interval). d) Overlap of TFa-target pairs from solid, hematological cell lines identified in a pan-cancer analysis. e) Overlap of TFa-target genes and the TF regulons used to estimate TF activity. Plots showing the distribution of Jaccard score of overlap between regulons and identified target genes across different types of analysis performed in this study.

**Supp Figure 2.**
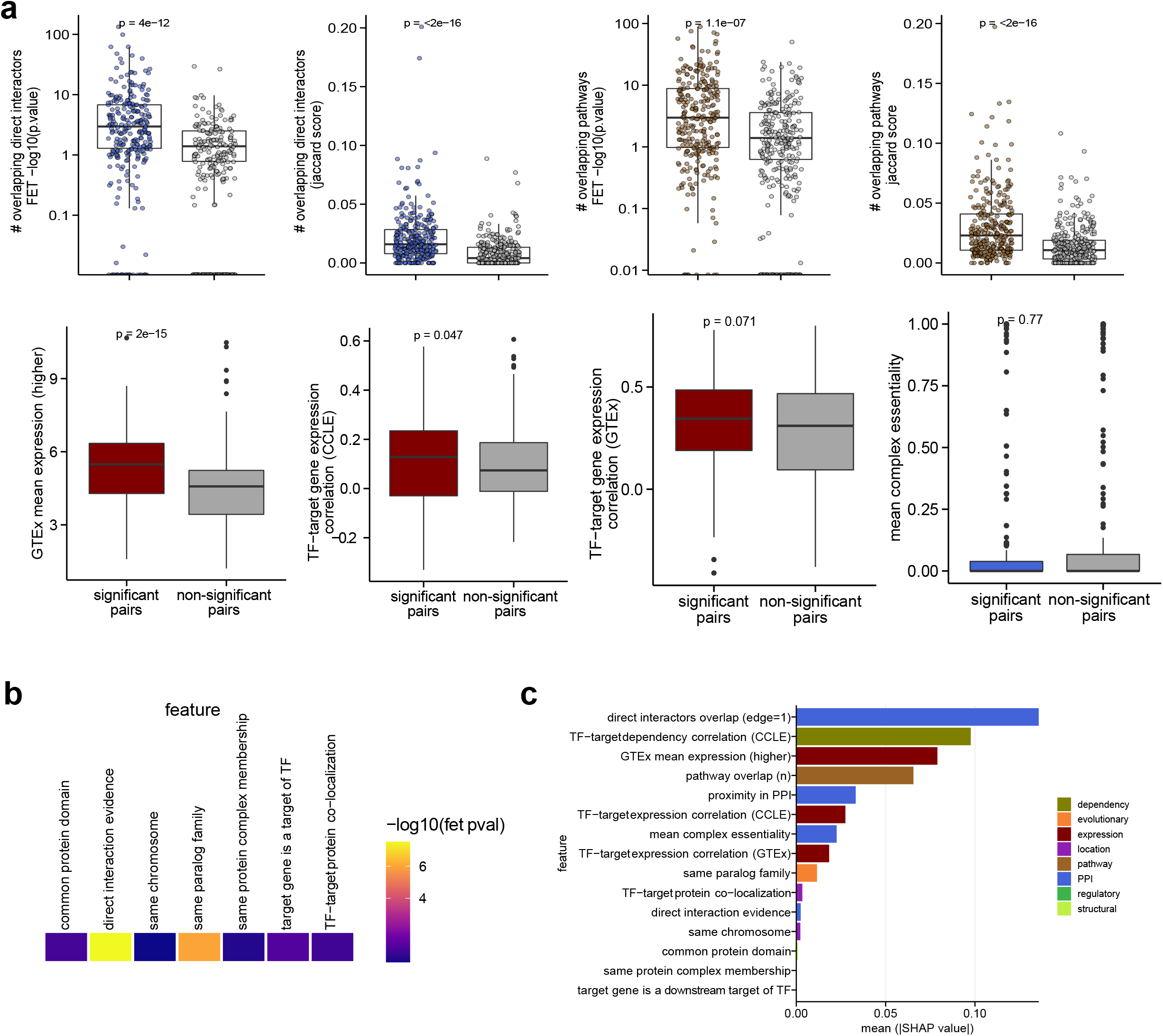
a) Box plots showing the difference of quantitative features between TFa-target significant vs non-significant pairs. b) P-values estimated by Fishers’ exact test for binary features. FET: Fisher’s exact test. c) SHAP values estimated for all features from random forest model are averaged across all pairs and ranked.

**Supp Figure 3.**
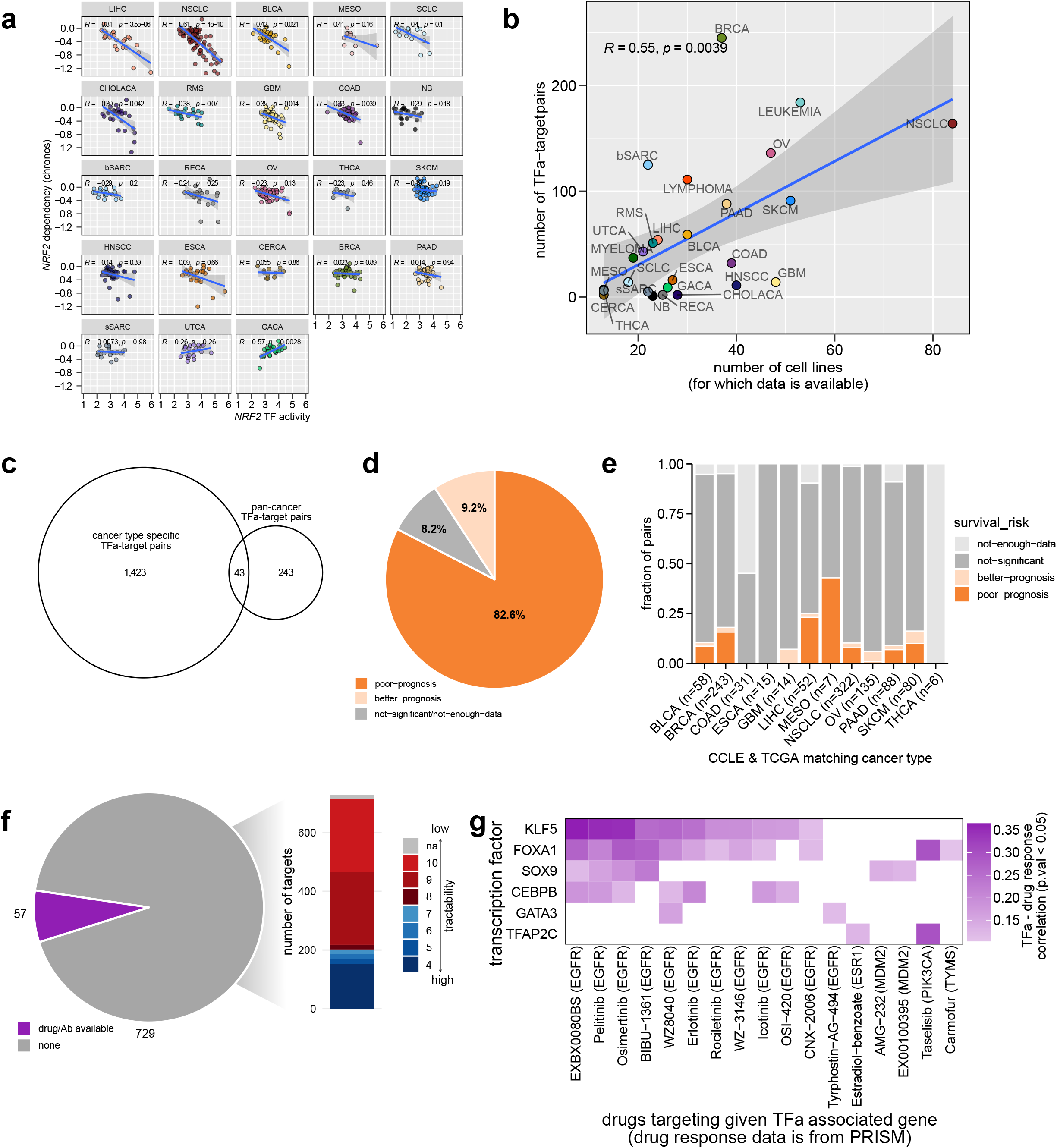
a) Scatter plot showing the correlation of NRF2 (also called NFE2L2) TFa vs its dependency across different cancer types. b) Scatter plot showing the correlation between the number of TFa-target pairs identified and the total number of cell lines available for each cancer type. c) Venn diagram showing the overlap of TFa-target pairs from pan-cancer analysis of solid cancer cell lines and pairs identified from cancer type specific analysis. d) Pie chart showing the distribution of TFa-target pairs identified by individual cancer type analysis and their association with survival in TCGA cancer types. e) Distribution of individual cancer type TFa-target pairs and their survival impact in matched TCGA cohorts. f) Pie chart indicates the number of target genes from analysis that currently have inhibitors and antibodies. The bar chart indicates the tractability buckets of the remaining 729 target genes. Lower the number for tractability bucket indicates the high tractability g Heatmap showing the correlation of inhibitor responses from PRISM database and TFa (spearman correlation, 95% confidence interval).

**Supp Figure 4.**
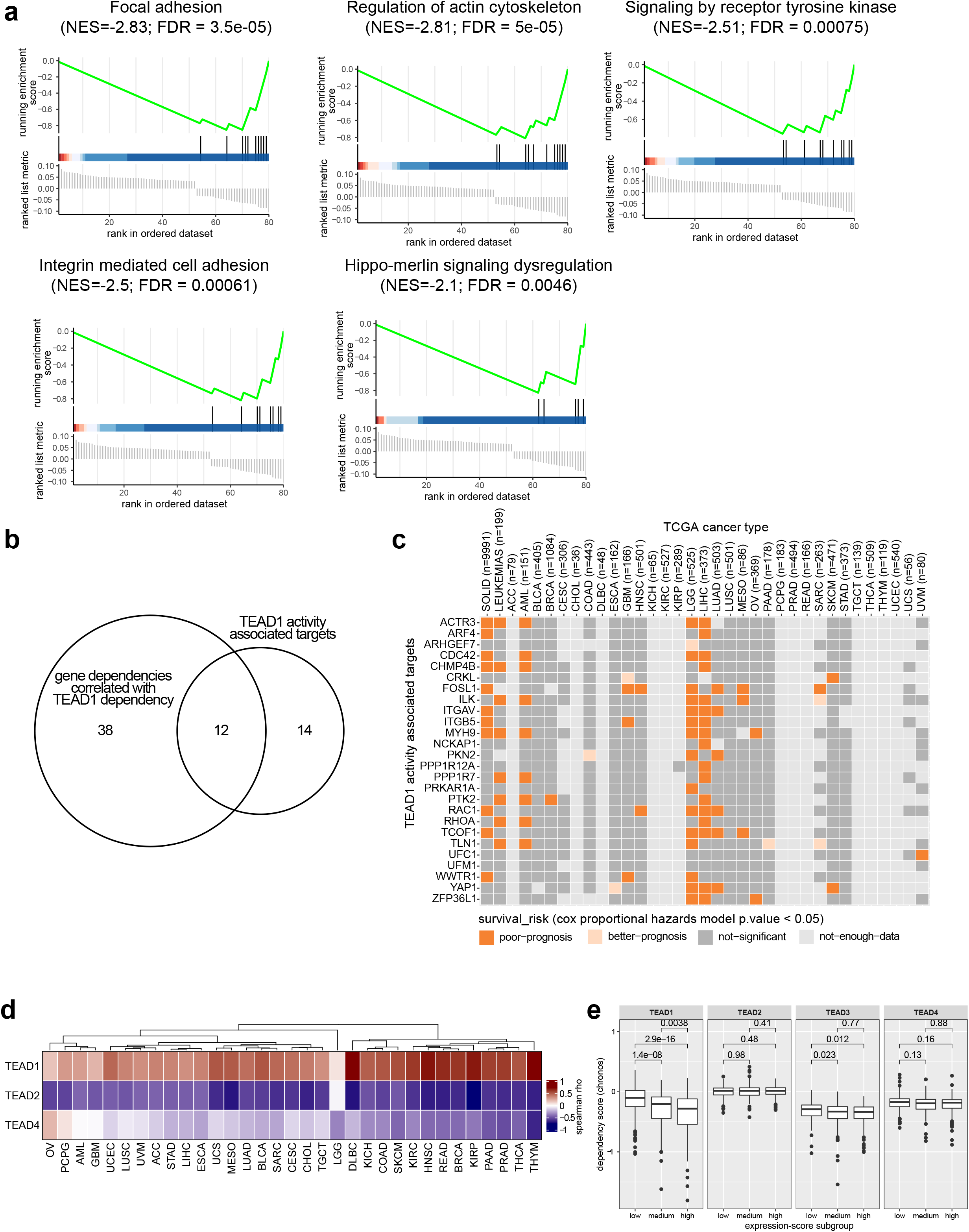
a) GSEA plots of enriched pathways with 26 target genes associated with TEAD1 TFa. b) Overlap of top 50 genes whose dependency is correlated with TEAD1 dependency and TEAD1 activity associated 26 targets genes. c) Heatmap showing the survival effect 26 target genes in TEAD1 activity high subgroup. TEAD1 activity high – low target expression subgroup is compared with TEAD1 activity high – high target expression subgroup. Both ‘poor-prognosis’ and ‘better-prognosis’ indicates p.value < 0.05 significance from cox proportional hazards model. d) Heatmap showing the correlation of expression-score of 26 genes as a gene set with different TEAD paralog activities in TCGA. Expression-score is estimated for each sample in TCGA using 26 target genes expression with single sample gene set enrichment analysis (spearman correlation, 95% confidence interval). e) Boxplots illustrating the dependency of TEAD paralogues in three subgroups defined based on 26 genes expression-score. 757 solid cancer cell lines were divided into three subgroups based on the expression-score of each cell line.

**Supp Figure 5.**
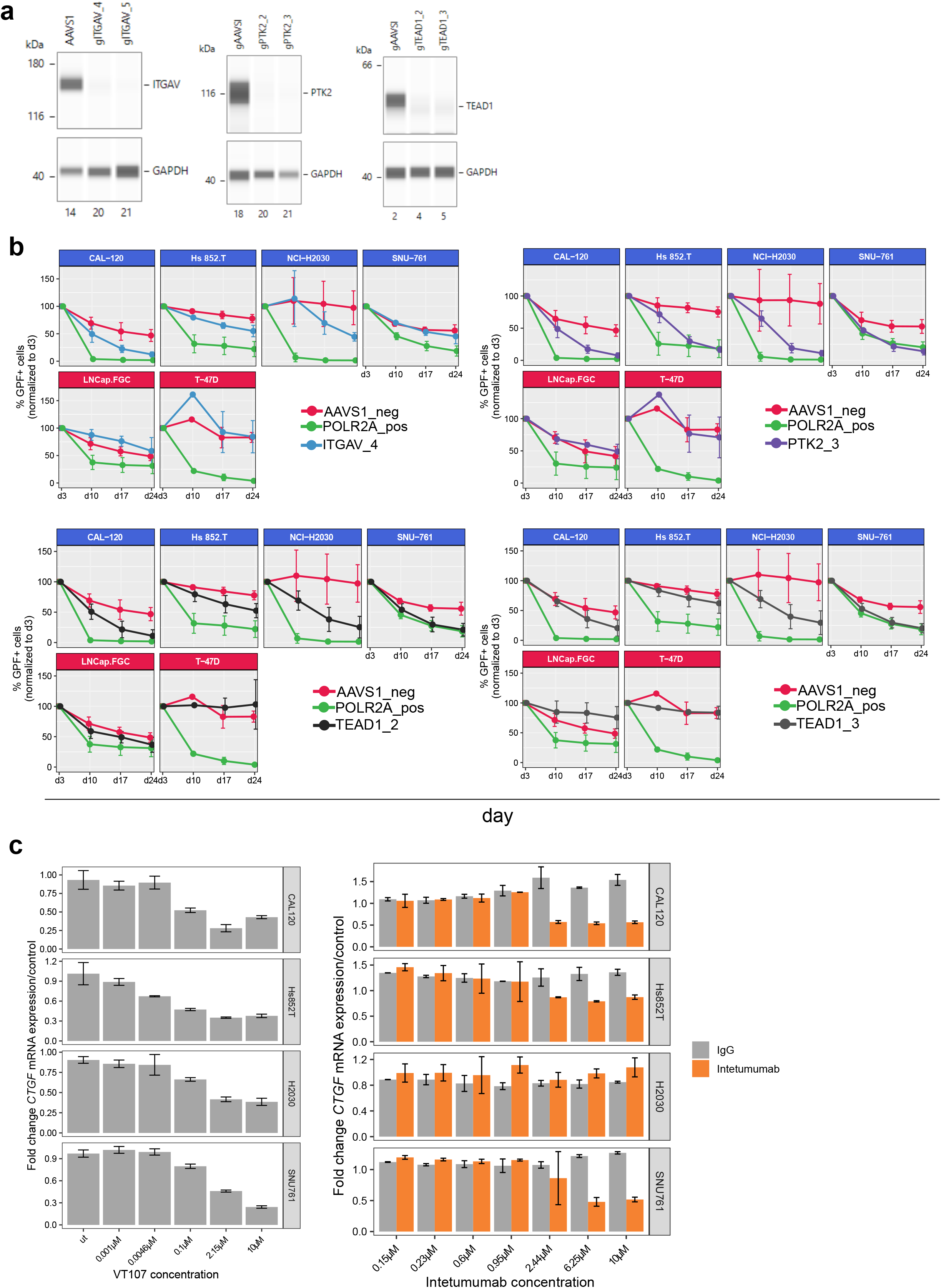
a) Automated Western blot (WES) showing reduced ITGAV protein levels in NCI -H2030 at d7 upon transduction with ITGAV-targeting guides. Reduced protein levels of PTK2 and TEAD1 in NCI-H2030 sorted for >95% purity at day 7 upon transduction with respective guides. b) CRISPR/Cas9 depletion time course showing relative growth defects introduced by sgRNAs against PTK2 (purple), ITGAV (blue), TEAD1 (black), the negative control AAVSI (red), or the positive control guide POLRA (green). Guide containing GFP+ ce **l** numbers of weekly measurementsover 24 days were normalized to the GFP+ cell numbers of day 3. Mean and standard deviation of three biological replicates are shown. For T-47D resistant cell line, two biological replicates were presented and with one measurement on day 10. c) TEAD1 downstream biomarker gene CTGF expression measured with qPCR after treatment with inhibitor VT107 (pan-TEAD inhibitor) and antibody Intetumumab (ITGAV antibody).

## Acknowledgements

We wish to thank Johannes Wachter, Nadja Liehmann, Eimilie Trebla, Dominik Arnold, Barbara Hopfgartner, Anju Kombara, Silvia Blaha-Ostermann and Alex Horman for technical assistance, David Schönbauer and Peter Greb for synthesis and QC of the VT103, VT107 and K975 compounds used in this study, and all colleagues at Boehringer Ingelheim RCV Cancer Research Target Discovery for discussions and critical input to the manuscript. We thank David Schönbauer and Peter Greb for performing analytical quality control of the compounds and Fiorella Schischlik-Siegl and Julie Livingstone for reading our manuscript and providing useful feedback.

## Author contributions

V. T, A.P conceptualized the study. Analyses were performed by V.T, V.S, F.M, A.P. Data visualization was conducted by V.T, V.S, F.M, A.P. Experiments were performed by V.S, F.M. A.S.B, M.G.R, M.M.R, J.A.R performed PRISM inhibitor screen. The original draft was written by V.T and A.P, and all co-authors reviewed and edited the manuscript.

